# Two opposing hippocampus to prefrontal cortex pathways for the control of approach and avoidance behavior

**DOI:** 10.1101/2019.12.18.880831

**Authors:** Candela Sánchez-Bellot, Andrew F. MacAskill

## Abstract

The decision to either approach or avoid a potentially threatening environment is thought to rely upon complex connectivity between heterogenous neural populations in the ventral hippocampus and prefrontal cortex (PFC). However, how this circuitry can flexibly promote both approach or avoidance at different times has remained elusive. Here, we show that the projection to PFC is composed of two parallel circuits located in the superficial or deep hippocampal pyramidal layers. These circuits have unique upstream and downstream connectivity, and are differentially active during approach and avoidance behavior. The superficial population is preferentially connected to widespread PFC inhibitory interneurons, and its activation promotes exploration; while the deep circuit is connected to PFC pyramidal neurons and fast spiking interneurons, and its activation promotes avoidance. Together this provides a mechanism for regulation of behavior during approach avoidance conflict: through two specialized, parallel circuits that allow bidirectional hippocampal control of PFC.

## INTRODUCTION

The decision to explore a novel or potentially threatening environment is essential for survival. Too little exploration reduces the likelihood of finding sources of reward, while too much increases risk such as injury. Dysfunction in such decision making, such as in the overestimation of threat, is thought to be a core feature of a number of anxiety disorders, and can lead to maladaptive avoidance behavior (Gray and McNaughton, 2003).

Resolving the conflict between approach and avoidance behavior is thought to crucially depend on the activity of the ventral hippocampus (vH) (Bannerman et al., 2003; Gray and McNaughton, 2003; Ito and Lee, 2016). Classically, vH activity is thought to inhibit approach behavior and thus increase avoidance during such conflicts. For example, in the elevated plus maze (EPM) - a commonly used assay of innate approach avoidance conflict - the decision to explore the innately threatening open arms, or to remain in the relative safety of the closed arms is thought to be defined by the overall level of vH activity (Bannerman et al., 2003; Gray and McNaughton, 2003; Jimenez et al., 2018; Kjelstrup et al., 2002). However, recent work has shown that vH activity can both promote or inhibit approach and avoidance behavior, dependent on how vH circuitry is manipulated (Jimenez et al., 2018; LeGates et al., 2018; Okuyama et al., 2016; Padilla-Coreano et al., 2016; 2019; Parfitt et al., 2017; Pi et al., 2020). Consistent with this more complex role, neuronal activity in vH during approach and avoidance is also heterogenous. In the EPM, different neurons in vH fire upon entry into either the open arms or the closed arms of the maze (Ciocchi et al., 2015; Jimenez et al., 2018), suggesting that different populations of neurons may be involved in the promotion of either approach or avoidance behavior.

The organization of the vH circuit is also heterogenous, and is composed of distinct, nonoverlapping subpopulations of neurons that vary in their morphology, physiology, and local and long-range connectivity (Cembrowski and Spruston, 2019; Soltesz and Losonczy, 2018). In particular, the downstream projection target of vH neurons is frequently used to distinguish functional specializations during behavior (Ciocchi et al., 2015; Jimenez et al., 2018; LeGates et al., 2018; Naber and Witter, 1998; Okuyama et al., 2016; Wee and MacAskill, 2020). Neurons in vH that preferentially fire during either approach or avoidance are both enriched in a subpopulation that project to the prefrontal cortex (PFC) (Ciocchi et al., 2015). Notably, the neurons that make up the vH-PFC projection are markedly heterogenous (Cembrowski et al., 2018), further suggesting that this projection may be made up of multiple populations of neurons with distinct functions. However, how this heterogeneity relates to function during behavior, and how this function is achieved via projections to PFC circuitry is unknown.

One way this may occur is via distinct subpopulations of neurons in vH differentially connecting to excitatory and inhibitory circuitry in PFC. Neurons in PFC track exploration in the EPM (Adhikari et al., 2011; Padilla-Coreano et al., 2016), and PFC activity defines behavior within the maze: increased excitatory activity in PFC increases avoidance behavior (Berg et al., 2019; Canetta et al., 2016; Soumier and Sibille, 2014), while PFC inhibition promotes approach (Green et al., 2020; Wall et al., 2004). Importantly, the extent of excitatory and inhibitory drive in PFC that results from hippocampal activation is different at unique points during behavior (Jadhav et al., 2016). Notably, during approach-avoidance conflict, the influence of vH input can be either inhibitory (Sotres-Bayon et al., 2012) or excitatory (Padilla-Coreano et al., 2016) dependent on ongoing behavior. Thus, the relative level of excitation and inhibition within PFC can be bidirectionally defined by vH, and is well placed to control the balance of approach and avoidance behavior. However, how heterogeneous populations of neurons in vH supports such flexible, bidirectional modulation of PFC is unknown.

Here we use a combination of *in vitro* and *in vivo* circuit analysis to show that the vH-PFC projection is made up of two populations of neurons distributed across the radial hippocampal axis. The activity of these two populations is controlled by specialized local and long-range afferent input, and each has unique connectivity in PFC. The superficially located population promotes exploration of the open arms of the EPM, and is preferentially connected to widespread inhibitory circuitry within PFC. The deep population promotes entry to the closed arms of the EPM, and is preferentially connected to excitatory pyramidal neurons and fast spiking interneurons. Together, our data support a model where two separate vH-PFC projections bidirectionally control behavior during approach avoidance conflict.

## RESULTS

### The hippocampal projection to prefrontal cortex consists of two populations distributed across the radial axis

To investigate the cellular organization of the projection from vH to PFC, we first labelled neurons that project to the PFC with an injection of the retrograde tracer cholera toxin (CTXβ, **Fig.1A**), and examined the distribution of fluorescently labelled neurons in transverse slices of vH. Strikingly, labelled neurons consistently formed two distinct layers at the CA1/subiculum border, located at the extreme poles of the radial axis (**Fig.1B**) that could reliably be separated with an unsupervised gaussian mixture model (**Sup.Fig.1**). This suggests that the vH-PFC projection is segregated along the radial axis of vH.

**Figure 1.**
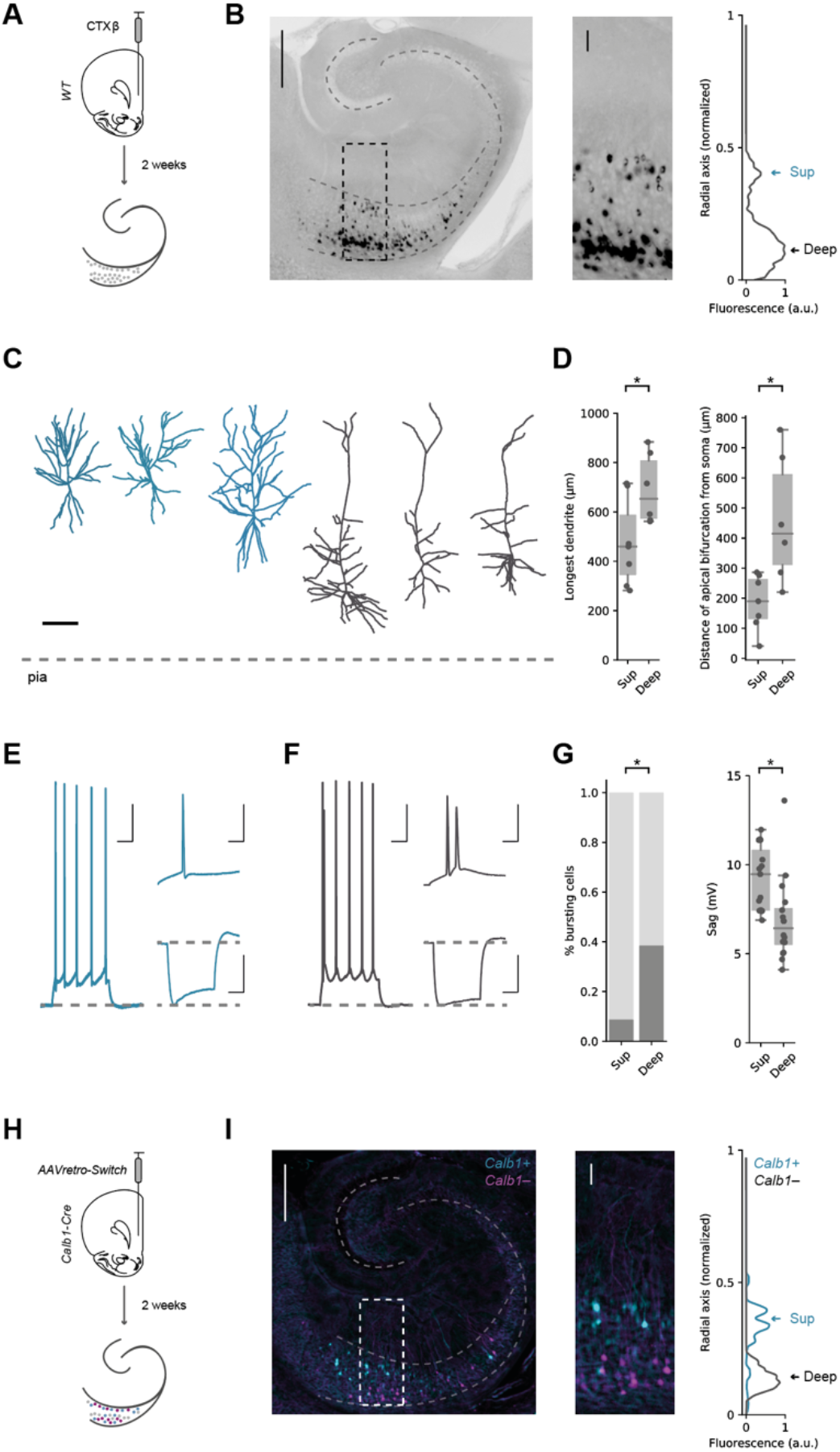
Hippocampal neurons projecting to PFC form two populations segregated across the radial axis. **A)** Schematic of cholera toxin (CTXβ) injection into PFC and retrograde labelling in hippocampus. **B)** *Left*, Transverse slice of hippocampus labelled with CTXβ. *Right*, zoom of retrogradely labelled neurons in boxed region, with fluorescence intensity profile. Arrows highlight the two distinct peaks of fluorescence at the two extremes of the radial axis. Scale bar = 500 μm (*Left*) 100 μm (*Right*). **C)** Example reconstructions of superficial (blue) and deep (dark grey) PFC-projecting hippocampal neurons. Scale bar = 100 μm, dotted line represents pia. **D)** *Right*, quantification of the distance of the apical bifurcation from the soma in superficial (Sup) and Deep neurons. *Left*, quantification of the distance from the farthest dendrite tip to the soma. **E)** *Left*, example response of a superficial layer PFC-projecting cell in response to a current injection of 140 pA. Top right, detail of first 50 ms of current injection. Bottom right, response to a current injection of −160 pA. Scale bars = 100 ms, 20 mV; 10 ms, 20 mV; 100 ms, 20 mV. **F)** As in (E) but for a neighboring deep layer PFC-projecting neuron. Note burst firing in response to current injection, and lower level of voltage sag after negative current injection. **G)** *Left*, quantification of the proportion of bursting neurons after positive current injections. *Right*, quantification of voltage sag. **H)** *AAVretro-Switch* injection into the PFC of *Calb1-Cre* mice and subsequent cre-dependent retrograde labelling in hippocampus. **I)** Transverse slice of hippocampus labelled with *AAVretro-Switch*. Cyan labels *Calb1^+^* PFC-projecting neurons and magenta labels *Calb1^-^* neurons. *Right*, zoom of retrogradely labelled neurons in boxed region, with fluorescence intensity profile for *Calb1^+^* (*cyan*) and *Calb1^-^* (*black*) neurons. Arrows highlight the two genetically distinct peaks of fluorescence at the two extremes of the radial axis. Scale bar = 500 μm (*Left*) 100 μm (*Right*). See **Sup.Fig.1** for further quantification.

To investigate if the two vH-PFC layers corresponded to previously identified circuits in superficial and deep layer hippocampus with specialized properties, we carried out targeted whole-cell current-clamp recordings in acute transverse slices of ventral hippocampus from mice previously injected with a retrograde tracer in PFC. By recording sequential pairs of retrogradely labelled neurons in the superficial and deep layers, this allowed us to compare intrinsic electrophysiological properties. In addition, filling the neurons with a morphological dye during the recording allowed us to reconstruct the dendritic morphology of a subset of these recorded neurons using two-photon microscopy. Superficial layer neurons were morphologically compact, with early branching of the apical dendrite, while deep layer neurons tended to have a relatively sparse, but long apical dendritic tree (**Fig.1C,D**). Electrophysiologically, superficial neurons were predominantly regular firing with high I_*h*_, while deep layer neurons were more likely to burst fire, and had a more subtle I_*h*_ (**Fig.1E-G**). These recordings further suggested that the vH-PFC projection is composed of two populations of neurons, each arising from the classical superficial and deep layer of the hippocampus (Cembrowski and Spruston, 2019; Harris et al., 2001; Jarsky et al., 2008; Soltesz and Losonczy, 2018).

We next utilized the expression of *Calb1* – a gene known to be specific to the superficial layer (Li et al., 2017; Pi et al., 2020), and injected a retrograde AAV expressing a fluorescence switch cassette into the PFC of a *Calb1-IRES-cre* mouse line. In this experiment, the color of the fluorophore expressed depended on the presence of *cre* (**Fig.1H**). Neurons projecting to PFC, that also express *Calb1* (and therefore *cre*) will be labeled with one fluorophore, while neurons projecting to PFC that do not express *Calb1* will be labeled with a different fluorophore (Saunders et al., 2012). The expression of *Calb1* robustly differentiated each population (**Fig.1I**), again suggesting that vH neurons that project to PFC can be split into two populations. Thus, the two vH-PFC populations have distinct molecular, cellular and physiological properties that are consistent with each arising from the superficial and deep layers of the hippocampus.

### Superficial and deep vH-PFC neurons are differentially connected to local and long-range input

vH is innervated by a wide range of afferent input, the identity of which is strongly influenced by both the spatial location and downstream projection of the neuron (Cembrowski and Spruston, 2019; Strange et al., 2014; Wee and MacAskill, 2020). Our results thus far suggest the two populations of vH-PFC neurons may be poised to receive different afferent input dependent on their downstream projection (to PFC), but also dependent on their spatial location along the radial axis (superficial or deep). To investigate if this was the case, we first used *tracing the relationship between input and output* (TRIO) – a rabies virus based retrograde tracing technique that allowed us to trace the input arriving specifically onto vH neurons projecting to PFC. Using TRIO we found dense input onto vH-PFC neurons from a number of local and long range regions (**Sup.Fig.2**).

We next used *channelrhodopsin assisted circuit mapping* (CRACM) to investigate the functional input to each layer (**Fig.2**). We used AAV injections to express channelrhodopsin (ChR2) using a pan-neuronal synapsin promoter in each of the 4 most densely labeled areas identified in our TRIO experiment – hippocampal CA3, medial entorhinal cortex (ENT), a disperse cluster of anterior thalamic regions focused around paraventricular thalamus (ATh) and an area encompassing the ventral medial septum and the diagonal band of Broca (DBB, see **Sup.Fig.2**). After 2 weeks to allow for virus expression, we made acute transverse slices of vH and performed sequential paired whole cell current-clamp recordings from retrogradely labelled PFC-projecting neurons in each layer. Using brief pulses of blue (473 nm) light allowed us to directly compare the relative synaptic input arriving into superficial or deep layer vH-PFC neurons from each of the afferent regions (MacAskill et al., 2014). We found that while CA3 terminal stimulation resulted in roughly equal excitatory synaptic drive in both superficial and deep neurons, ENT input was biased towards superficial layer neurons, and both DBB and ATh input was markedly biased towards deep layer neurons (**Fig.2A-D, Sup.Fig.2**). Thus, the two populations of vH-PFC neurons are differentially connected to afferent input – both populations receive dense CA3 input, but superficial cells receive additional input from cortex, while deep cells receive additional predominantly thalamic and basal forebrain input.

**Figure 2.**
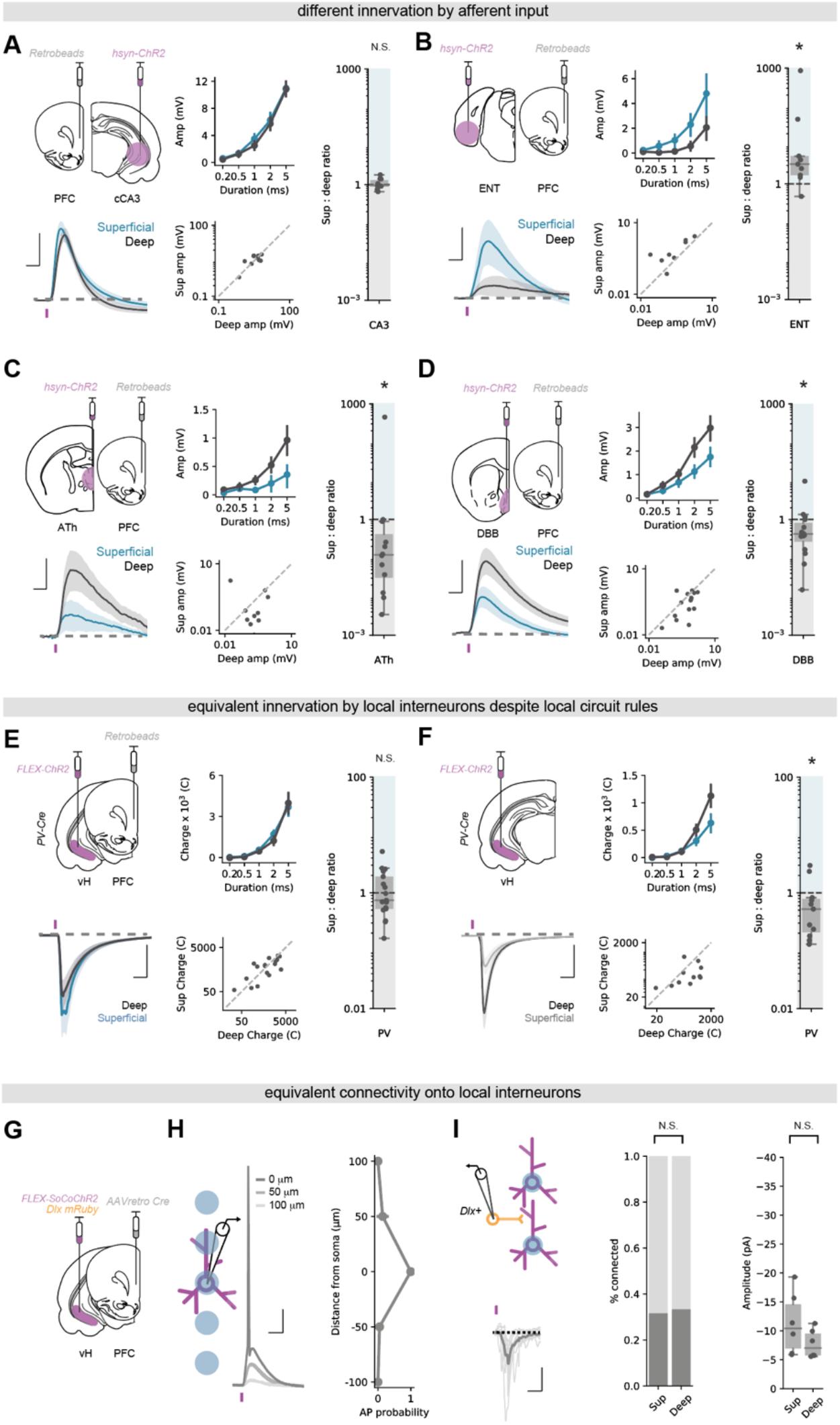
Superficial and deep vH-PFC neurons are differentially connected to local and long-range input. **A)** *Top left*, Schematic showing experimental setup. ChR2 was injected into contralateral CA3 and retrobeads injected into PFC. 2 weeks later input-specific connectivity was assessed using paired recordings of superficial and deep vH-PFC neurons in acute slices. *Bottom left*, Average light-evoked responses in pairs of superficial (*blue*) and deep (*black*) layer PFC-projecting hippocampal neurons in response to cCA3 input. Scale bar = 10 ms, 5 mV. Purple tick represents the light stimulus. *Middle*, summary of amplitude of Sup and Deep responses to increasing durations of light pulse (*top*), and amplitudes of individual pairs at 5 ms (*bottom*). *Right*, summary of the ratio of superficial : deep neuron EPSP. Higher values mean input is biased to superficial neurons, low values towards deep layer neurons. Note log scale. CA3 input is equivalent onto superficial and deep layer neurons. **B-D)** As in **(A)** but for ENT **(B)**, ATh **(C)** and DBB **(D)** input. Scale bar = 10 ms, 2 mV (**B**), 0.5 mV (**C**), 1 mV (**D**). ENT input is biased towards superficial layer neurons, while both ATh and DBB are biased towards deep layer neurons. **E)** As in **(A)** but for local PV interneuron input. Scale bar = 10 ms, 500 pA. PV+ inhibitory input is equivalent onto superficial and deep layer PFC-projecting neurons. **F)** As in **(E)** but in neighboring, unlabeled vH neurons from superficial and deep layers. Scale bar = 10 ms, 200 pA. PV+ inhibitory input is biased towards non-retrogradely labeled deep layer neurons. **G)** Strategy to investigate superficial and deep vH neuron connectivity onto local interneurons. **H)** Focused light allows activation of neurons expressing soCoChR with high spatial resolution. Scale bar = 10 mV, 20 ms. **I)** Connectivity of superficial and deep PFC projecting vH neurons onto neighboring dlx+ interneurons. Probability of connectivity and amplitude is equivalent for both layers. Scale bar = 10 pA, 20 ms.

Excitatory local connectivity in hippocampal CA1 and subiculum is rare, however, there is strong local inhibition mediated by interneurons (Cembrowski and Spruston, 2019; Lee et al., 2014). In particular, the connectivity of parvalbumin positive (PV+) interneurons in vH is dependent on both spatial location along the radial axis, and downstream projection target. Therefore, we next investigated how the two populations of vH-PFC neurons were connected with the local interneuron network.

To investigate the connectivity of local PV+ interneurons onto each layer, we used AAV injections in a *PV-IRES-Cre* mouse to express ChR2 in PV+ interneurons in vH, and retrograde tracers to label PFC-projecting neurons (**Fig.2E**). After 2 weeks we performed sequential paired whole-cell voltage-clamp recordings from each layer, using a high chloride internal to allow for isolation of inhibitory currents. Similar to before, we could then use brief blue light pulses to investigate PV mediated inhibition onto superficial and deep vH-PFC neurons. We found that ChR2-mediated activation of PV+ interneurons in vH resulted in robust, yet equivalent IPSCs in both superficial and deep layer vH-PFC neurons (**Fig.2E**). This was surprising, as it is in contrast to previous reports of preferential inhibition of deep layer neurons (Lee et al., 2014). Therefore, in the same slices we recorded neighboring unlabeled superficial and deep neurons, and confirmed that non-PFC-projecting vH neurons in the deep layers receive more input than superficial layers (**Fig.2F**). Thus, vH-PFC projecting neurons are specifically connected to ensure equivalent inhibition across the two layers.

Next, to investigate the excitatory input onto local interneurons arising from vH-PFC neurons in each layer, we combined an injection of retrograde AAV expressing cre recombinase into PFC, and an injection into vH of a mixture of AAV to express cre-dependent fluorescently tagged somatic channelrhodopsin (soCoChR2, Shemesh et al., 2017), and the fluorescent marker mRuby under the control of the interneuron specific *dlx* promoter (Cho et al., 2015; Dimidschstein et al., 2016). This allowed us to elicit action potentials in individual PFC-projecting neurons from each layer using a focused light spot to activate somatically targeted CoChR2, while recording from neighboring genetically identified interneurons in vH (**Fig.2G,H**). Using this approach, we again found that PFC-projecting neurons in each layer connected to local interneurons with a similar connection probability and synaptic strength (**Fig.2I**). Combined with the disparate afferent input identified above, this suggests that the two vH-PFC circuits are connected in a way that might facilitate lateral inhibition across the two layers, to promote their activity at distinct timepoints.

### Superficial and deep vH-PFC neurons connect differentially in PFC

Our results so far suggest the presence of two populations of vH-PFC neurons that are differentially controlled by local and long-range input. We next asked if these populations might connect differentially onto excitatory and inhibitory neurons in PFC.

We first used monosynaptic rabies tracing from either inhibitory interneurons (using *VGAT+ starter cells*) or excitatory neurons (using *CaMKii+ starter cells*) in PFC (**Fig.3A**). We compared neurons retrogradely labelled with rabies in vH, with the distribution of neurons labelled with a simultaneous injection of CTXβ in PFC. Therefore, in this experiment all vH-PFC neurons are labelled with CTXβ, while either vH-PFC^excitatory^ or vH-PFC^inhibitory^ neurons are labelled with rabies. Using this strategy, we found differences in the targeting of excitatory and inhibitory neurons in PFC by each layer in vH (**Fig.3B**). While, both superficial and deep layer vH neurons targeted interneurons in PFC, input to pyramidal neurons in PFC preferentially originated from neurons in deep layers.

**Figure 3.**
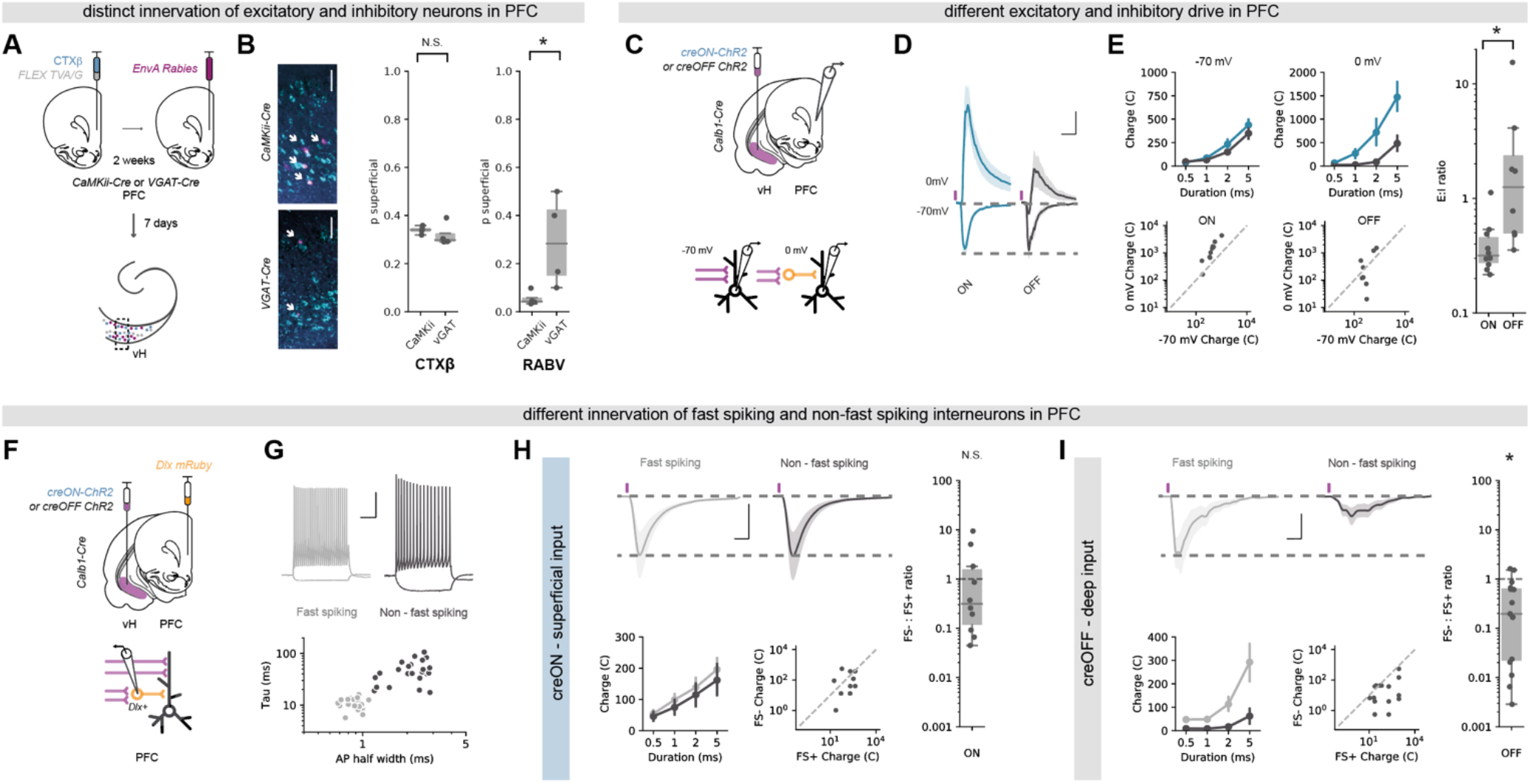
Superficial and deep vH-PFC neurons connect differentially in PFC. **A)** Strategy to label neurons projecting to inhibitory and excitatory neurons in PFC. **B)** *Left*, Transverse slice of hippocampus labelled with CTXβ (cyan) and rabies (magenta) after tracing from excitatory (top) or inhibitory (bottom) neurons in PFC. Note the restriction of rabies labelling from excitatory neurons to the deep layer. Scale bar = 100 μm. *Right*, Proportion of CTXβ positive neurons (left) and rabies positive neurons (right) in the superficial layer. Note equivalent distribution of CTXβ across both conditions, but a marked absence of neurons projecting to excitatory PFC neurons in the superficial layer. **C)** Strategy to record E:I ratio in PFC from each layer in vH. **D)** Responses to superficial (blue) or deep (grey) hippocampal inputs at −70 mV (EPSCs) and 0 mV (IPSCs) in deep layer PFC neurons. Purple tick indicates light pulse. Scale bar = 20 ms and 0.5 (fold response amplitude at −70 mV, which is normalized to 1). **E)** *Left*, Summary of amplitude of superficial (creON, blue) and deep (creOFF, grey) responses at −70 mV and 0 mV to increasing durations of light pulse (*top*), and amplitudes of individual responses for superficial, and deep input at 5 ms (*bottom*). *Right*, summary of the ratio of −70 mV : 0 mV. Higher values mean input is biased to excitation, low values towards inhibition. Note log scale. Input from superficial neurons has a greater inhibitory contribution than that of deep neurons. **F)** Strategy to record input from the two layers of vH onto identified interneurons in PFC. **G)** *Top*, example current clamp recordings from fast-spiking (FS+, grey) and non-fast-spiking (FS-, black) neurons in PFC. *Bottom*, summary showing clustering into two groups based on membrane time constant and action potential half width. **H)** *Top*, responses to superficial (creON) input at −70 mV (EPSCs) onto neighboring FS (grey) or non-FS interneurons (black). Scale bar = 10 ms, 200 pA. *Bottom*, summary of amplitude of FS+ and FS− responses to increasing durations of light pulse (*left*), and amplitudes of individual pairs at 5 ms (*right*). *Right*, summary of the ratio of FS− : FS+. Higher values mean input is biased to FS-, low values towards FS+ neurons. Note log scale. Superficial input is equivalent onto FS+ and FS− neurons. **I)** As in **(H)** but for deep (creOFF) input. Deep input is biased towards FS+ interneurons.

Our tracing experiment suggested that the superficial layers connect more readily with inhibitory interneurons in PFC, while deep layer neurons connect with a mixture of both inhibition and excitation. To confirm this distinct targeting using CRACM, we used *Calb1-IRES-cre* mice, and injected an AAV into vH to express either ChR2 where expression was limited to *cre*-expressing (superficial layer) neurons (creON ChR2), or where ChR2 was inhibited in *cre*-expressing neurons, and thus was only expressed in *cre*-negative (deep layer) neurons (creOFF ChR2, **Fig.3C**, Saunders et al., 2012). We then carried out whole-cell voltage-clamp recordings of deep layer pyramidal neurons in acute slices of PFC from these animals in the presence of the NMDA-receptor antagonist APV, and assessed the relative excitatory and inhibitory drive by recording light-evoked synaptic input at −70 mV (predominantly AMPA-mediated excitatory currents) and 0 mV (predominantly GABA-mediated inhibitory currents, **Fig.3D**). By comparing the ratio of responses at −70 mV and 0 mV, we confirmed that the superficial layer of vH drives substantially more feedforward inhibition in PFC compared to the deep layer (**Fig.3E**).

Interneurons in PFC can be characterized into two main subgroups – soma-targeting fast-spiking interneurons, and dendrite-targeting non-fast-spiking interneurons – based on their intrinsic properties and the expression of peptides such as parvalbumin and somatostatin (The Petilla Interneuron Nomenclature Group (PING), 2008). As both deep and superficial layer vH neurons could drive feedforward inhibition - albeit to different extents - we next wanted to see if the inhibitory circuitry each layer contacted was different. To do this we again used CRACM to compare the relative input from superficial and deep layer vH neurons into PFC. However, we used an injection of AAV to express *dlx* mRuby into PFC to allow targeted whole cell recordings from inhibitory interneurons (**Fig.3F**). Using this approach, we classified each interneuron as either fast spiking or non-fast spiking based on intrinsic electrophysiological properties (**Fig.3G**), and then examined the relative input onto neighboring pairs of neurons of each type from either the superficial or deep layer of vH. Superficial vH input innervated both fast-spiking and non-fast-spiking interneurons in PFC to an equal extent, suggesting widespread recruitment of both dendritic and somatic inhibitory circuits within PFC (**Fig.3H**). In contrast, deep layer vH input was very selective - while it showed reliable input onto fast-spiking interneurons in PFC, there was little to no input onto neighboring non-fast-spiking interneurons (**Fig.3I**). Together this suggests that vH-PFC neurons are connected in such a way to have differential influence on PFC.

### Superficial and deep vH-PFC neurons have opposing activity around open and closed arm entry in the EPM

The two populations of vH-PFC neurons are connected in a way that may support opposing activity, thus, we next asked if the superficial and deep populations in vH were differentially active during approach avoidance behavior.

To do this, we simultaneously recorded the activity of both populations using bulk calcium imaging through an optical fiber (**Fig.4A,B**). We used an injection of a mixture of retrograde AAVs into PFC to express both creON RGeCO1a – a red wavelength calcium indicator, and creOFF GCaMP6f – a green wavelength calcium indicator - in *Calb1-IRES-cre* mice. Using this technique, *Calb1+*, superficial neurons in vH projecting to PFC will express RGeCO1a, and *Calb1*− deep neurons projecting to PFC will express GCaMP6f. By collecting green and red fluorescence through an implanted optical fiber, this allowed simultaneous monitoring of both superficial and deep layer vH-PFC neuron activity in freely behaving mice while they explore the EPM.

**Figure 4.**
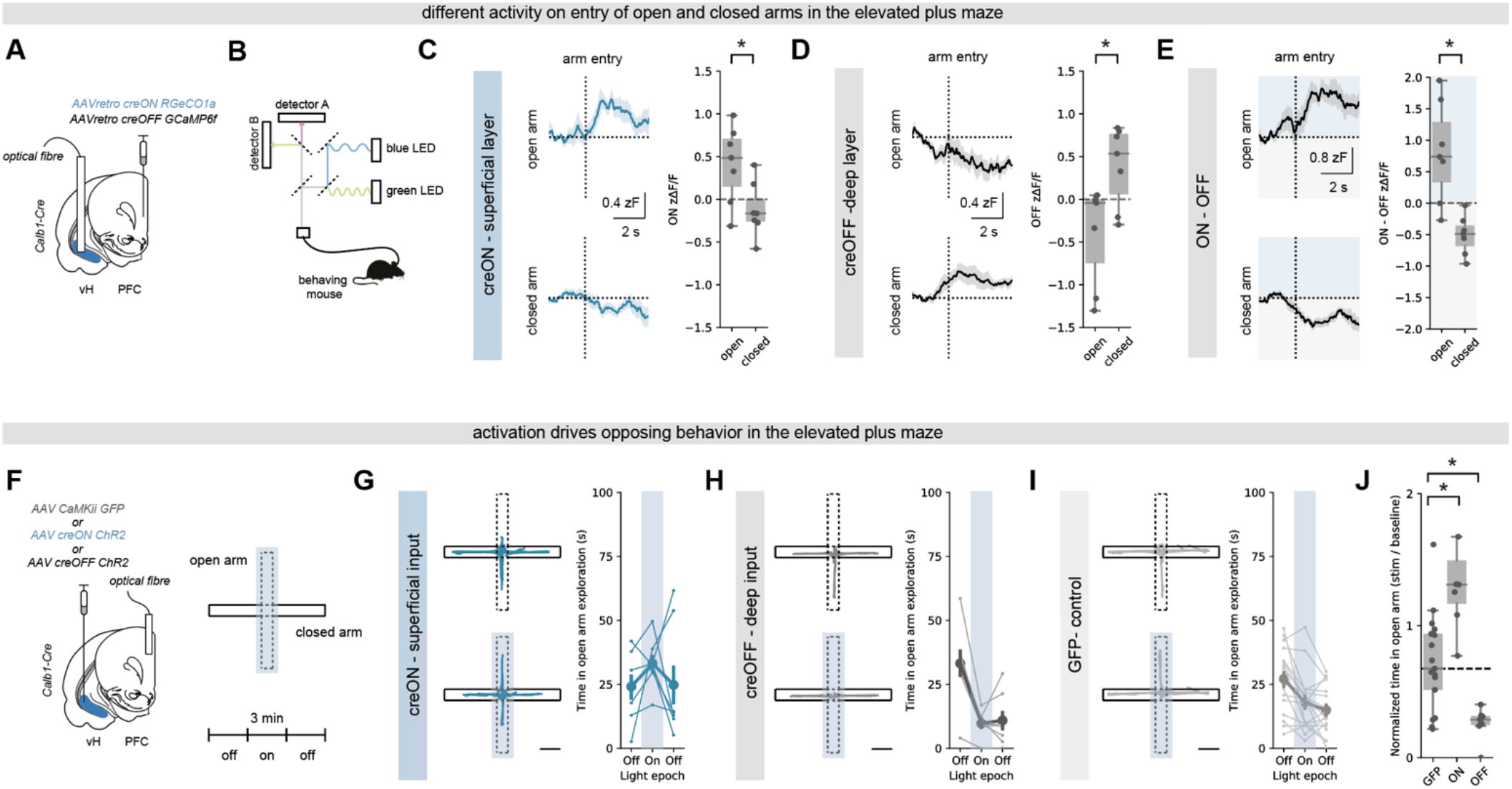
Superficial and deep vH-PFC populations bidirectionally influence behavior in the EPM. **A,B)** Strategy to record superficial and deep vH-PFC neuron calcium dynamics during free behavior. **C)** *Left*, superficial layer (creON) calcium fluorescence aligned to open arm (*top*) and closed arm (*bottom*) entry. Superficial neurons increase activity on open arm entry, and slightly decrease on closed arm entry. Right, summary showing superficial activity is greater on open arm entry compared to closed arm entry. **D)** As for **(C)** but for deep layer (creOFF) fluorescence. In contrast to superficial layer neurons, deep layer neurons decrease activity in response to open arm entry, and increase activity upon closed arm entry. **E)** As for **(C)** but for the difference between simultaneously recorded superficial (creON) and deep (creOFF) layer activity. **F)** *Left*, strategy for *in vivo* optogenetic manipulation of vH axons from each layer in PFC. *Right*, experimental design: after a 3 min baseline, for a second 3 min epoch 20 Hz light was delivered via the optical fiber when the mouse entered the center point of the maze, and continued until return to the closed arms. Mice then remained in the maze for a third post stimulation 3 min epoch. **G)** Superficial (creON) stimulation in PFC. *Left*, trajectories of an example mouse during baseline (top) and during stimulation (bottom). *Right*, change in open arm exploration due to stimulation. Scale bar = 10 cm. **H-I)** As in **(G)** but for deep (creOFF, **H**), or control (GFP, **I**) animals. **J)** Summary of the effect of activation on open arm exploration. Superficial (creON) stimulation increased, while deep (creOFF) decreased exploration relative to controls. Dotted line shows median exploration of GFP controls for comparison.

We aligned these recordings to when mice made the decision to move from the closed arms to the open arm of the EPM, and compared this to the equivalent decision to enter the closed arms of the maze. We found that upon exploration of the open arms of the maze, superficial neurons increased their activity. In contrast, upon entry to the closed arms of the maze, superficial neurons decreased activity (**Fig.4C**). Deep layer neurons showed the opposing pattern of activity at both these behavioral epochs. Activity of deep layer neurons was reduced upon entry of the open arms, and increased upon entry of the closed arms (**Fig.4D**). This suggested that the relative activity of the superficial and deep layers of the vH-PFC projection around the choice point of the EPM may inform the decision to approach or avoid the open arms (**Fig.4E**).

### Optogenetic activation of superficial or deep input into PFC has opposing influence on behavior in the EPM

Finally, we wanted to test the causal role of the two vH-PFC populations in the exploration of the EPM. Due to the large differences in connectivity and utilization during behavior, we hypothesized that artificial activation of the superficial and deep layers may drive opposite behavior.

We again used an AAV based approach in *Calb1-IRES-cre* mice to express ChR2, or control GFP, in either superficial (creON) or deep (creOFF) layer neurons in vH, and implanted optical fibers unilaterally in PFC (**Fig.4F**). This allowed us to stimulate axons arriving in PFC from each of the two layers of vH with blue light while the animal was exploring the maze. We artificially activated axons with blue light for a three-minute epoch, only when mice entered the central choice point in the EPM, and maintained excitation until the mice returned to the closed arms. We found that this manipulation consistently increased open arm exploration in creON (superficial layer) mice (**Fig.4G**), while in contrast dramatically reduced open arm exploration in creOFF (deep layer) mice (**Fig.4H**), compared to GFP controls (**Fig.4I**). Therefore, activation of superficial layer vH axons in PFC promotes the exploration of the open arms of the EPM, while activating the deep layer vH axons in PFC reduces exploration of the open arms (**Fig.4J**).

## DISCUSSION

Here we show that the projection from vH to PFC can be subdivided into two populations that are segregated along the radial axis of vH (**Fig.1,2**). These parallel populations of pyramidal neurons have opposing influence on both PFC circuit activity (**Fig.3**) and behavior during innate approach avoidance conflict (**Fig.4**). The superficially located population is preferentially connected to widespread inhibitory circuitry in PFC, is driven by cortical input and promotes exploration. In contrast, the deep population is preferentially connected to pyramidal neurons and fast spiking interneurons in PFC, is driven by basal forebrain and thalamic input, and promotes avoidance.

We used a combination of whole brain anatomy and CRACM to investigate the connectivity of these two populations of neurons. Superficial vH-PFC neurons receive strong input from cortex (**Fig.2B**, (Li et al., 2017; Masurkar et al., 2017), consistent with a role for superficially located neurons in the relay of stable information, and utilization during prospective decision making (Danielson et al., 2016; Mizuseki et al., 2011; Valero et al., 2015). Distinct from superficial neurons in dorsal hippocampus, which receive preferential input from lateral entorhinal cortex (Li et al., 2017), input onto vH-PFC neurons derives from medial areas. This is consistent with medial entorhinal cortex providing the predominant input to the ventral location occupied by vH-PFC neurons (Canto et al., 2008). Together this suggests superficial neurons may use structured information from cortex to plan and promote exploratory, approach behavior (**Fig.4**) (Buzsáki and Moser, 2013). In contrast, deep layer vH-PFC neurons receive preferential input from subcortical areas such as anterior thalamus and the diagonal band (**Fig.2C,D**) which are associated with value, salience and attention, consistent with a role in flexible updating (Danielson et al., 2016; Soltesz and Losonczy, 2018, Jimenez et al., 2018). Input from these areas is strongly theta modulated (Soltesz and Losonczy, 2018), consistent with more prominent theta modulation of deep layer neurons (Mizuseki et al., 2011). Notably, neurons in PFC that preferentially encode anxiogenic environments are strongly coupled to hippocampal theta (Adhikari et al., 2011; 2010; Lee et al., 2019). Together this suggests that deep layer vH-PFC neurons may use salient information in the environment flexibly to promote avoidance (**Fig.4**).

Consistent with these proposed roles, superficial vH-PFC neurons preferentially connect with wide feedforward inhibition in PFC, including strong input onto dendrite-targeting non-fast spiking interneurons (**Fig.3**). Consistent with the effect of superficial layer activation in the EPM (**Fig.4**), dendrite targeting interneurons have been implicated in ongoing utilization of spatial memory, and the promotion of approach and exploration (Abbas et al., 2018; Soumier and Sibille, 2014). In contrast, deep layer vH input preferentially connects directly with pyramidal neurons and with fast-spiking interneurons, which have been implicated in flexible updating, and the promotion of avoidance (Berg et al., 2019; Canetta et al., 2016; Marek et al., 2018), again consistent with the effect of deep layer vH-PFC activation in the EPM (**Fig.4**). Together this suggests a key role for hippocampal input from each layer in dynamically controlling the utilization of excitatory and inhibitory PFC circuitry to control approach and avoidance behavior.

With this in mind, disruption in the balance of excitation and inhibition in PFC is closely associated with the transition to mental illness (Gao and Penzes, 2015). Alterations in vH input to PFC are thought to be key for this disruption, in particular in response to chronic stress and genetic mutations associated with schizophrenia (Canetta et al., 2016; Mukherjee et al., 2019; Sigurdsson et al., 2010). Our study reveals a mechanism by which changes in the activity of each layer in vH could exert strong shifts in excitatory and inhibitory balance in PFC. Thus, understanding how these two layers may be differentially impacted in models of mental illness is an interesting future avenue of investigation.

Overall, our findings provide a mechanism for the regulation of behavior during approach avoidance conflict: through two specialized parallel circuits that allow bidirectional hippocampal control of PFC.

## ACKNOWLEDGEMENTS

We thank Marco Tripodi for providing rabies virus, and Francesca Cacucci for help with surgery. We thank Neil Burgess, Tara Keck, Thomas Mrsic-Flogel, Aman Saleem, Marcus Stephenson-Jones, Tom Wills, and members of the MacAskill laboratory for helpful comments on the manuscript. We also thank David Attwell, Francesca Cacucci, Mark Farrant, Troy Margrie and Andreas Schaefer for comments on a previous version. A.F.M. was supported by a Sir Henry Dale Fellowship jointly funded by the Wellcome Trust and the Royal Society (grant number 109360/Z/15/Z) and by a UCL Excellence Fellowship. C.S.B. was supported by the Wellcome Trust 4-year PhD in Neuroscience at UCL (grant number 206074/Z/17/Z).

## AUTHOR CONTRIBUTIONS

Conceptualization, C.S.B. and A.F.M.; Methodology, C.S.B. and A.F.M.; Investigation, C.S.B. and A.F.M.; Formal Analysis, C.S.B. and A.F.M.; Writing – Original Draft, C.S.B. and A.F.M.; Writing – Review & Editing, C.S.B. and A.F.M.; Funding Acquisition, C.S.B. and A.F.M.; Supervision, A.F.M.

## DECLARATION OF INTERESTS

The authors declare no competing interests.

## DATA AVAILABILITY

The data that support the findings of this study are available from the corresponding author upon reasonable request.

## METHODS

### Animals

6 - 10 week old (adult) male and female C57 / bl6J mice provided by Charles River were used except where noted. To target inhibitory neurons we used the *Slc32a1(VGAT)-IRES-Cre* (#016962) knock-in line. To target Calbindin expressing neurons we used the *Calb1-IRES-Cre* (#028532) knock-in line. To target parvalbumin positive interneurons we used the *PV-IRES-Cre* (#008069) knock in line. All were obtained from Jackson laboratories and bred in-house. Mice were housed in cages of 2 - 4 and kept in a humidity- and temperature-controlled environment under a 12 h light/dark cycle (lights on 7 am to 7 pm) with ad-libitum access to food and water. All experiments were approved by the U.K. Home Office as defined by the Animals (Scientific Procedures) Act, and University College London ethical guidelines.

### Stereotaxic surgery

#### Retrograde tracers

Red and Green fluorescent retrobeads (Lumafluor, Inc.) for electrophysiological recordings. Cholera toxin subunit B (CTXβ) tagged with Alexa 555 or 647 (Molecular Probes) for histology experiments.

#### Viruses

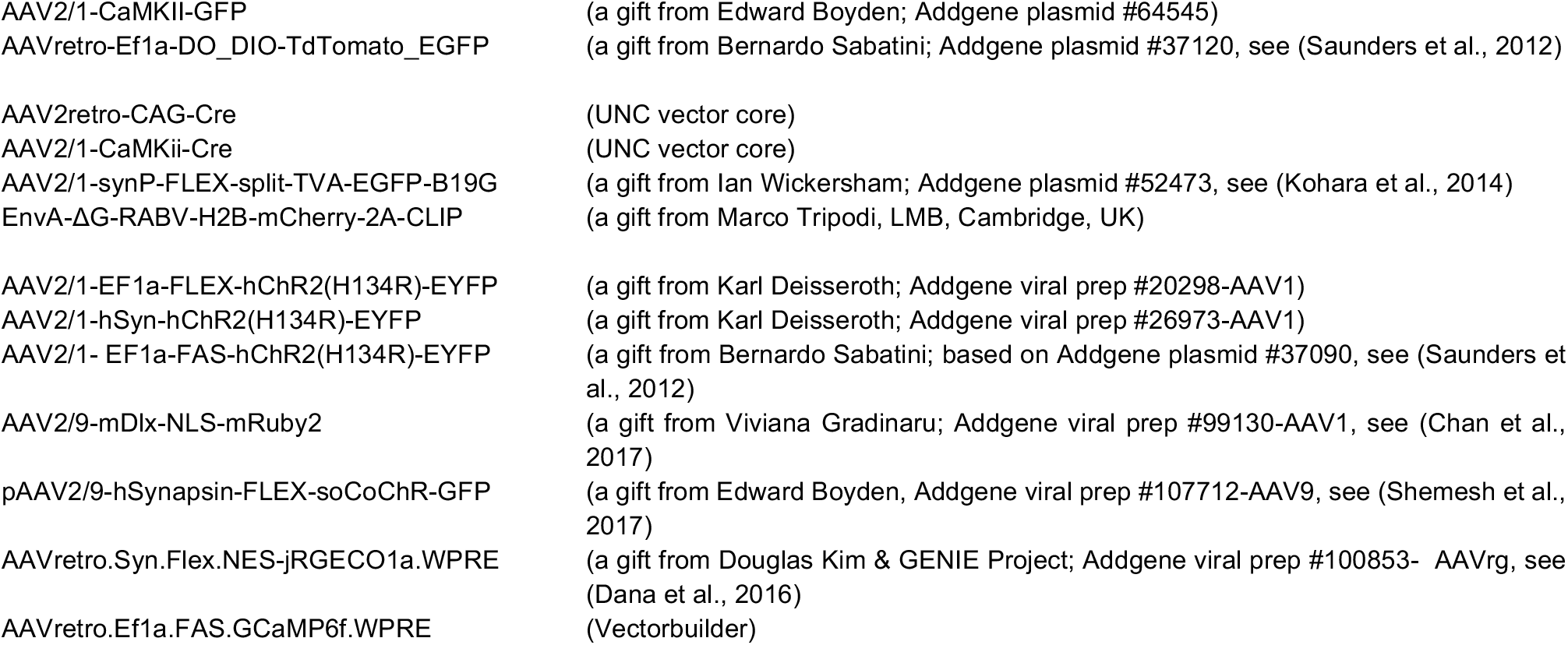

#### Surgery

Stereotaxic injections were performed on 7 - 10 week old mice anaesthetized with isofluorane (4 % induction, 1 - 2 % maintenance) and injections carried out as previously described (Little and Carter, 2013; MacAskill et al., 2014; 2012). Briefly, the skull was exposed with a single incision, and small holes drilled in the skull directly above the injection site. Injections are carried out using long-shaft borosilicate glass pipettes with a tip diameter of ~ 10 - 50 μm. Pipettes were back-filled with mineral oil and front-filled with ~ 0.8 μl of the substance to be injected. A total volume of 140 - 160 nl of each virus was injected at each location in ~ 14 or 28 nl increments every 30 s. If two or more substances were injected in the same region they were mixed prior to injection. The pipette was left in place for an additional 10 min to minimize diffusion and then slowly removed. If optic fibers were also implanted, these were inserted immediately after virus injection, secured with 1 – 2 skull screws and cemented in place with C&B superbond. Injection coordinates were as follows (mm relative to bregma):

|  |  |  |  |  |
| --- | --- | --- | --- | --- |
| 1 |  |  |  |  |
| 2 | infralimbic PFC: | ML: $\pm 0.4$, | RC: + 2.3, | and DV: - 2.4, |
| 3 | Diagonal band of Broca: | ML: $\pm 0.2$, | RC: + 0.7, | and DV: - 4.0, |
| 4 | anterior thalamus: | ML: $\pm 0.2$, | RC: - 0.4, | and DV: - 3.2, |
| 5 | CA3: | ML: $\pm 2.8$, | RC: - 3.0, | and DV: - 4, |
| 6 | Entorhinal cortex: | ML: $\pm 2.7$, | RC: - 5.2 (@ 10 °), | and DV: - 4.5, |
| 7 | ventral hippocampus: | ML: $\pm 3.4$, | RC: - 3.7, | and DV: - 4.3. |
| 8 |  |  |  |  |

After injection, the wound was sutured and sealed, and mice recovered for ~30 min on a heat pad before they were returned to their home cage. Animals received carprofen in their drinking water (0.05 mg / ml) for 48 hrs post-surgery as well as subcutaneously following surgery (0.5 mg / kg). Expression occurred in the injected brain region for ~2 weeks until preparation of acute slices for physiology experiments, or fixation for histology. The locations of injection sites were verified for each experiment.

### Anatomy

#### Pseudotyped rabies labelling from PFC

For pseudotyped rabies experiments two injections were necessary. First, *AAV2/1-synP-FLEX-split-TVA-EGFP-B19G* was injected into the PFC of either *VGAT-Cre* mice to express TVA and G protein in inhibitory neurons, or was coinjected in a 1:1 mixture with *AAV2/1-CaMKii-Cre* into Cre negative littermates to label excitatory neurons. In addition, CTXβ was coinjected into PFC to achieve non-specific retrograde labelling for comparison across mice. 2 weeks later, 200 nl of *EnvA-ΔG-RABV-H2B-mCherry* was injected into PFC to infect and induce transynaptic spread only in neurons expressing Cre. Animals were prepared for histology after 7 days of rabies-mediated expression.

#### TRIO labelling from PFC-projecting vH neurons

TRIO labelling (Schwarz et al., 2015) again required two surgeries. First, *AAVretro-CAG-Cre* was injected into the PFC to express Cre recombinase in neurons projecting to PFC. In the same surgery *AAV2/1-synP-FLEX-split-TVA-EGFP-B19G* was injected into the vH to express rabies TVA and G-protein only in vH neurons that express Cre (i.e. only those that project to PFC). 2 weeks later, 200 nl of *EnvA-ΔG-RABV-H2B-mCherry* was injected into vH to infect and induce trans-synaptic spread only in PFC-projecting vH neurons. Animals were prepared for histology after 7 days of rabies-mediated expression.

#### Histology

Mice were perfused with 4% PFA (wt / vol) in PBS, pH 7.4, and the brains dissected and postfixed overnight at 4°C as previously described (MacAskill et al., 2012; 2014). 70 μm thick slices were cut using a vibratome (Campden Instruments) in either the transverse, coronal or sagittal planes as described in the figure legends. Slices were mounted on Superfrost glass slides with ProLong Gold or ProLong Glass (for visualization of GFP) antifade mounting medium (Molecular Probes). NucBlue was included to label gross anatomy. Imaging was carried out with a Zeiss Axio Scan Z1, using standard filter sets for excitation/emission at 365-445/50 nm, 470/40-525/50 nm, 545/25-605/70 nm and 640/30-690/50 nm. Raw images were analyzed with FIJI.

#### Analysis of spatial distribution of PFC-projecting hippocampal neurons

The spatial distribution of retrogradely labelled PFC-projecting hippocampal neurons was analyzed in transverse slices from mice injected with cholera toxin in PFC, and imaged as described above. 3 slices spanning 300μm between ~−3.5 and −5.0 mm DV were analyzed per injection. Sections containing the hippocampus were straightened along the cell body layer using the straighten macro in FIJI to produce a single field stretching from proximal CA3 to distal subiculum. Labelled cells within each slice were manually counted, and the coordinates saved for later analysis. The majority of cells were found at the CA1 / subiculum border - defined as the point where the hippocampal cell layer widens into a more subicular-like structure, which occurs consistently at ~0.7 of the distance of the entire straightened field. Custom scripts written in Python based around the scikit.learn package were used to cluster the coordinates using a Gaussian Mixture Model (GMM). Models containing between 1 and 6 components were fit to the data, and Bayesian Information Criterion (BIC) was used to select the best fit.

#### Analysis of rabies labelling from VGAT and CaMKii neurons in PFC

As above, transverse hippocampal slices containing cholera toxin and rabies labelling of PFC-projecting cells were used for cell counting. All cells in the straightened hippocampal formation were counted, irrespective of fluorescent label to gain an overall distribution of vH-PFC neurons. These coordinates were clustered using GMM as above, where again, all analyzed fields were best fit by two components. Rabies positive cells were assigned to one of the two components (i.e. superficial and deep layers) created by the clustering algorithm. Data is presented as proportion of all rabies labelled cells in each layer. As an internal control data for cholera toxin labelling is also presented using the same method, which shows there are no differences in the overall layer structure between the *CaMKii* and *VGAT* experiments.

### Electrophysiology

#### Slice preparation

Hippocampal recordings were studied in acute transverse slices, while prefrontal cortical recordings in **Fig.3** were studied in acute coronal slices. Mice were anaesthetized with a lethal dose of ketamine and xylazine, and perfused intracardially with ice-cold external solution containing (in mM): 190 sucrose, 25 glucose, 10 NaCl, 25 NaHCO_3_, 1.2 NaH_2_PO_4_, 2.5 KCl, 1 Na^+^ ascorbate, 2 Na^+^ pyruvate, 7 MgCl_2_ and 0.5 CaCl_2_, bubbled with 95% O_2_ and 5% CO_2_. Slices (300 μm thick) were cut in this solution and then transferred to artificial cerebrospinal fluid (aCSF) containing (in mM): 125 NaCl, 22.5 glucose, 25 NaHCO_3_, 1.25 NaH_2_PO_4_, 2.5 KCl, 1 Na^+^ ascorbate, 3 Na^+^ pyruvate, 1 MgCl_2_ and 2 CaCl_2_, bubbled with 95% O_2_ and 5% CO_2_. After 30 min at 35 °C, slices were stored for 30 min at 24 °C. All experiments were conducted at room temperature (22–24 °C). All chemicals were from Sigma, Hello Bio or Tocris.

#### Whole-cell electrophysiology

Whole-cell recordings were made from hippocampal pyramidal neurons retrogradely labelled with retrobeads or CTXβ, which were identified by their fluorescent cell bodies and targeted with Dodt contrast microscopy, as previously described (Little and Carter, 2013; MacAskill et al., 2012; 2014). For sequential paired recordings, neurons were identified within a single field of view at the same depth into the slice. The recording order was counterbalanced to avoid any potential complications that could be associated with rundown. For current clamp recordings, borosilicate recording pipettes (2– 3 MΩ) were filled with (in mM): 135 K-gluconate, 10 HEPES, 7 KCl, 10 Na-phosphocreatine, 10 EGTA, 4 MgATP, 0.4 NaGTP. For voltage clamp experiments, three internals were used, First, in **Fig.2I** and **Fig.3D,E**, a Cs-gluconate based internal was used containing (in mM): 135 Gluconic acid, 10 HEPES, 7 KCl, 10 Na-phosphocreatine, 4 MgATP, 0.4 NaGTP, 10 TEA and 2 QX-314. In **Fig.2I** excitatory currents were isolated by recording at − 70 mV in 10 mM extracellular Gabazine. In **Fig.3D,E**, excitatory and inhibitory currents were electrically isolated by setting the holding potential at −70 mV (excitation) and 0 mV (inhibition) and recording in the presence of 10 mM extracellular APV. Second, to allow characterization of interneuron intrinsic properties, experiments in **Fig.3F-I** were carried out using current clamp internal and excitatory currents were isolated using 10 mM extracellular Gabazine. Finally, to record inhibitory currents at rest in **Fig.2E,F**, we used a high chloride internal (in mM): 135 CsCl, 10 HEPES, 7 KCl, 10 Na-phosphocreatine, 10 EGTA, 4 MgATP, 0.3 NaGTP, 10 TEA and 2 QX-314, with 10 mM external NBQX and 10 mM external APV. For two-photon experiments, the internal solution also contained 30 μM Alexa Fluor 594 (Molecular probes). Recordings were made using a Multiclamp 700B amplifier, with electrical signals filtered at 4 kHz and sampled at 10 kHz.

#### Viral strategy for in vitro optogenetics

Presynaptic glutamate release was triggered by illuminating ChR2 in the presynaptic terminals of long-range inputs into the slice, as previously described (Little and Carter, 2013; MacAskill et al., 2012; 2014). Wide-field illumination was achieved via a 40 × objective with brief (0.2, 0.5, 1, 2 and 5 ms) pulses of blue light from an LED centered at 473 nm (CoolLED pE-4000, with appropriate excitation-emission filters). Light intensity was measured as 4–7 mW at the back aperture of the objective and was constant between all cell pairs.

To achieve afferent specific terminal stimulation in vH (**Fig.2A-D**), AAV2/1-hSyn-hChR2(H134R)-EYFP was injected into each downstream area, and retrobeads were injected into PFC to label the two layers in vH.

To achieve input-specific terminal stimulation in PFC (**Fig.3**), we injected AAV2/1-EF1a-FLEX-hChR2(H134R)-EYFP (creON) or AAV2/1-EF1a-FAS-hChR2(H134R)-EYFP (creOFF) into *Calb1-IRES-Cre* mice to target superficial neurons, and deep layer neurons respectively.

To target whole cell recordings to interneurons (**Fig.2I, Fig.3F-I**), we used AAV2/9-dlx-mRuby injections in the area of interest.

To express ChR2 in local PV interneurons within vH (**Fig.2E,F**), we injected AAV2/1-EF1a-FLEX-hChR2(H134R)-EYFP into the vH of *PV-IRES-Cre* mice, and retrobeads into PFC to allow paired recordings from the two layers.

To allow single cell optogenetic stimulation (**Fig.2G-I**), we used an injection of AAVretro-syn-Cre in PFC, and an injection of AAV2/1-EF1a-DIO-SoCoChR in vH. The resultant sparse labelling of neurons across each layer allowed stimulation of individual SoCoChR expressing neurons with a focused light spot (with an activation resolution of ~50 μm **Fig.2H**), while recording from neighboring interneurons labelled with dlx-mRuby as above.

#### 2 photon imaging and image reconstruction

Two-photon imaging was performed with a customized Slicescope (Scientifica), based on a design previously described (Little and Carter, 2013; MacAskill et al., 2012; 2014). For two-photon imaging, 780 nm light from an Erbium fiber laser (Menlo Systems) was used to excite Alexa Fluor 594. For each experiment a high-resolution stack of the entire neuron was taken for reconstruction of dendrite morphology.

#### Electrophysiology data acquisition and analysis

Two-photon imaging and physiology data were acquired using National Instruments boards and SciScan (Scientifica) and WinWCP (University of Strathclyde) respectively. Electrical stimulation was via a tungsten bipolar electrode (WPI) and an isolated constant current stimulator (Digitimer). Optical stimulation was via wide field irradiance with 473 nm LED light (CoolLED) as described above. Data was analyzed using custom routines written in Python 3.6. Current step data was analyzed using routines based around the Neo and eFEL packages. For synaptic connectivity experiments, amplitudes of PSPs were taken as averages over a 2-ms time window around the peak. For connectivity analysis, a cell was considered connected if the light-induced response was greater than 6 times the standard deviation of baseline.

Two-photon image reconstruction and analysis was carried out using VIAS and NeuronStudio (Dumitriu et al., 2011), before further analysis was carried out using custom scripts in Python 3.6 based on the NeuroML package from the Human Brain Project. The online Human Brain Project morphology viewer was used for visualizing reconstructed neurons.

### Behavior

After sufficient time for surgical recovery and viral expression (>4 weeks), mice underwent multiple rounds of habituation. Mice were habituated to the behavioral testing area in their home cage for 30 min prior to testing each day. Mice were habituated to handling for at least 3 days, followed by 1-2 days of habituation to the optical tether in their home cage for 10 min.

#### Bulk calcium imaging using fiber photometry during elevated plus maze exploration

##### Labelling superficial and deep vH neurons for photometry

To allow simultaneous recordings of both superficial and deep vH-PFC populations, AAVretro-DIO-RGeCO1a (creON) and AAVretro-FAS-GCaMP6f (creOFF) were co-injected into PFC of *Calb1-IRES-Cre* mice as a 50:50 mix, and a 2.5 mm ferrule containing a 200 μm optical fiber was implanted in vH. This strategy allowed dual color imaging, as the red sensor RGeCO1a is expressed in superficial neurons, while the green sensor GCaMP6f is expressed in deep neurons in vH. FAS was used to counteract known interference between different Cre-dependent viruses (Saunders et al., 2012).

##### Bulk calcium imaging during behavior

After habituation (above), mice were exposed to the elevated plus maze (EPM) for 9 min, and allowed to freely explore the open and closed arms of the maze. To record calcium activity, we used a system based on (Kim et al., 2016; Lerner et al., 2015; Markowitz et al., 2018). Briefly, two excitation LEDs (565 nm ‘green’ and 470nm ‘blue’) were controlled via a custom script written in Labview (National Instruments). Blue excitation was sinusoidally modulated at 210Hz and passed through a 470 nm excitation filter while green excitation was modulated at 500 Hz and passed through a 565 nm bandpass filter. Excitation light from each LED was collimated, then combined using a 520 nm long-pass dichroic mirror. The excitation light was coupled into a high-NA (0.53), low-autofluorescence 200 μm patch cord by reflection with a multiband dichroic mirror and fiber coupler. Each LED was set to 100 μW at the far end of the patch cord, which was terminated with a 2.5 mm ferrule. The emission signal was collected through the same patch cord and collimator, and separated from the excitation light by the multiband dichroic. Green and red signals were split using a longpass dichroic mirror before being passed through a GFP emission filter and RFP emission filter respectively. Filtered emission was then collected by a femtowatt photoreceiver (Newport). The photoreceiver signal was sampled at 100 kHz, and each of the two modulated signals generated by the two LEDs was independently recovered using standard synchronous demodulation techniques implemented in real time by a custom Labview script. Three separated signals; green, red and autofluorescence (the signal from each sensor at the modulation frequency that does not match the sensor), were then downsampled to 50 Hz before being exported for further analysis. A least-squares linear fit was applied to the autofluorescence signal to align it to the green and red signals. Time series data was then calculated for each behavioral session as the zscore of (green or red signal - fitted autofluorescence signal). Calcium signals around open versus closed arm entries were then calculated by aligning normalized traces to timestamps from automated video analysis (Deeplabcut, Mathis et al., 2018), where traces were aligned to the moment the head entered the open arms. Activity was analyzed as the area under the curve from 0 to 4 s after arm entry, normalized to the area under the curve −4 to −2 s before arm entry. All optical components were from Thorlabs, Semrock or Chroma, patch cords were from Doric Lenses.

#### Optogenetic manipulation of neurons during elevated plus maze exploration

Axon terminals in PFC from superficial and deep vH neurons were labelled using a creON and creOFF approach as described above (see ***Optogenetics***), and a 200 μm optical fiber was implanted unilaterally above right PFC. Mice expressing GFP in vH were used as controls. After habituation (above), behavior was assessed using the EPM. As above, EPM sessions lasted 9 min. For stimulation, the session was split into 3, 3 min epochs. 20 Hz light stimulation was delivered via a 473 nm laser, coupled to a patch cord (6-8 mW at the end of the patch cord) during the second three-minute epoch to activate ChR2 positive vH terminals in PFC. Real-time light delivery was based on the location of the mouse in the EPM apparatus, where light stimulation occurred only during points where the mouse was in the center or open arms of the maze (Jimenez et al., 2018; Lee et al., 2019). Time spent investigating the open arms of the maze was scored for each mouse in each epoch using automated analysis (Deeplabcut, Mathis et al., 2018). Open arm exploration was defined as entry of the head into the open arm, so as to include stretch attend, prospective exploration. To avoid ceiling effects in optogenetic behavioral experiments, mice were excluded if they failed to enter either open arm in the first three minutes of testing. This exclusion criterion was pre-established before the start of the experiment (see Adhikari et al., 2015).

### Statistics

Summary data are reported throughout the figures either as boxplots, which show the median, 75^th^ and 95^th^ percentile as bar, box and whiskers respectively, or as line plots showing mean +/− s.e.m. Example physiology and imaging traces are represented as the mean +/− s.e.m across experiments. Data were assessed using statistical tests described in the supplementary statistics summary. Significance was defined as *P* < 0.05, all tests were two sided. No statistical test was run to determine sample size a priori. The sample sizes we chose are similar to those used in previous publications. Animals were randomly assigned to a virus cohort (e.g. ChR2 versus GFP), and where possible the experimenter was blinded to each mouse’s virus assignment when the experiment was performed. This was sometimes not possible due to e.g. the presence of the injection site in the recorded slice.

## STATISTICAL TEST SUMMARY

1 STATISTICAL TEST SUMMARY
| FIGURE | DESCRIPTORS | n | TEST USED | STATISTIC | P-VALUE |
| --- | --- | --- | --- | --- | --- |
| 1D | Longest dendrite (µm) | Sup: 7<br>Deep: 6 | Mann Whitney | U = 35.5 | p = 0.045 |
|  | Distance of bifurcation (µm) | Sup: 7<br>Deep: 6 | Mann Whitney | U = 38.5 | p = 0.015 |
| 1G | % bursting neurons | Sup: 22<br>Deep: 26 | Fishers Exact | Odds ratio = 12.5 | p = 0.009 |
|  | Sag amplitude (mV) | Sup: 22<br>Deep: 26 | Mann Whitney | U = 39.5 | p = 0.002 |
| 2A | CA3 sup : deep amp at 5 ms (mV) | 10 | Wilcoxon Paired | W = 27 | p = 1.0 |
| 2B | ENT sup : deep amp at 5 ms (mV) | 9 | Wilcoxon Paired | W = 44 | p = 0.007 |
| 2B | ATh sup : deep amp at 5 ms (mV) | 7 | Wilcoxon Paired | W = 12 | p = 0.034 |
| 2B | ENT sup : deep amp at 5 ms (mV) | 15 | Wilcoxon Paired | W = 24 | p = 0.041 |
| 2E | PV PFC+ sup:deep charge at 5 ms (C) | 17 | Wilcoxon Paired | W = 57 | p = 0.38 |
| 2F | PV PFC- sup:deep charge at 5 ms (C) | 11 | Wilcoxon Paired | W = 10 | p = 0.042 |
| 2I | % connected | Sup: 19<br>Deep: 18 | Fishers Exact | Odds ratio = 0.95 | p = 1.0 |
|  | Amplitude of connection if present (pA) | Sup: 5<br>Deep: 5 | Mann Whitney | U = 8 | p = 0.42 |
| 3B | CTX *p* superficial | Sup: 4<br>Deep: 4 | Mann Whitney | U = 4 | p = 0.3 |
|  | Rabies *p* superficial | Sup: 4<br>Deep: 4 | Mann Whitney | U = 16 | p = 0.03 |
| 3E | creON vs creOFF<br>-70 mV charge at 5 ms (C) | creON: 10<br>creOFF: 8 | Mann Whitney | U = 50 | p = 0.41 |
|  | creON vs creOFF<br>0 mV charge at 5 ms (C) | creON: 10<br>creOFF: 8 | Mann Whitney | U = 91 | p = 0.01 |
|  | creON vs creOFF<br>E : I ratio at 5 ms | creON: 10<br>creOFF: 8 | Mann Whitney | U = 10 | p = 0.006 |
| 3H | creON FS-: FS+ charge at 5 ms (C) | 10 | Wilcoxon Paired | W = 35 | p = 0.49 |
| 3H | creOFF FS-: FS+ charge at 5 ms (C) | 15 | Wilcoxon Paired | W = 111 | p = 0.02 |
| 4C,D | creON vs creOFF<br>zΔF/F open vs closed arm | 7 | Repeated measures<br>ANOVA<br>Main effect of layer:<br>Interaction between<br>layer and arm | F(1,6) = 20.0<br>F(1,6) = 20.0 | p = 0.004<br>p = 0.004 |
| 4C | creON<br>zΔF/F open vs closed arm | 7 | Wilcoxon Paired | W = 27 | p = 0.031 |
| 4D | creOFF<br>zΔF/F open vs closed arm | 7 | Wilcoxon Paired | W = 1 | p = 0.031 |
| 4E | creON – creOFF<br>zΔF/F open vs closed arm | 7 | Wilcoxon Paired | W = 27 | p = 0.031 |
| 4G-I | Time exploring open arm (s) | GFP: 17<br>creON: 7<br>creOFF: 7 | Repeated measures<br>ANOVA<br>Main effect of<br>epoch:<br>Interaction between<br>group and epoch: | F(2, 27) = 6.25<br>F(4, 56) = 4.86 | p = 0.006<br>p = 0.002 |
| 4J | Normalised open arm exploration | GFP: 17<br>creON: 7 | Mann Whitney | U = 12 | p = 0.002 |
|  | Normalised open arm exploration | GFP: 17<br>creOFF: 7 | Mann Whitney | U = 100 | p = 0.008 |

## SUPPLEMENTARY FIGURES

**Sup. Fig 1.**
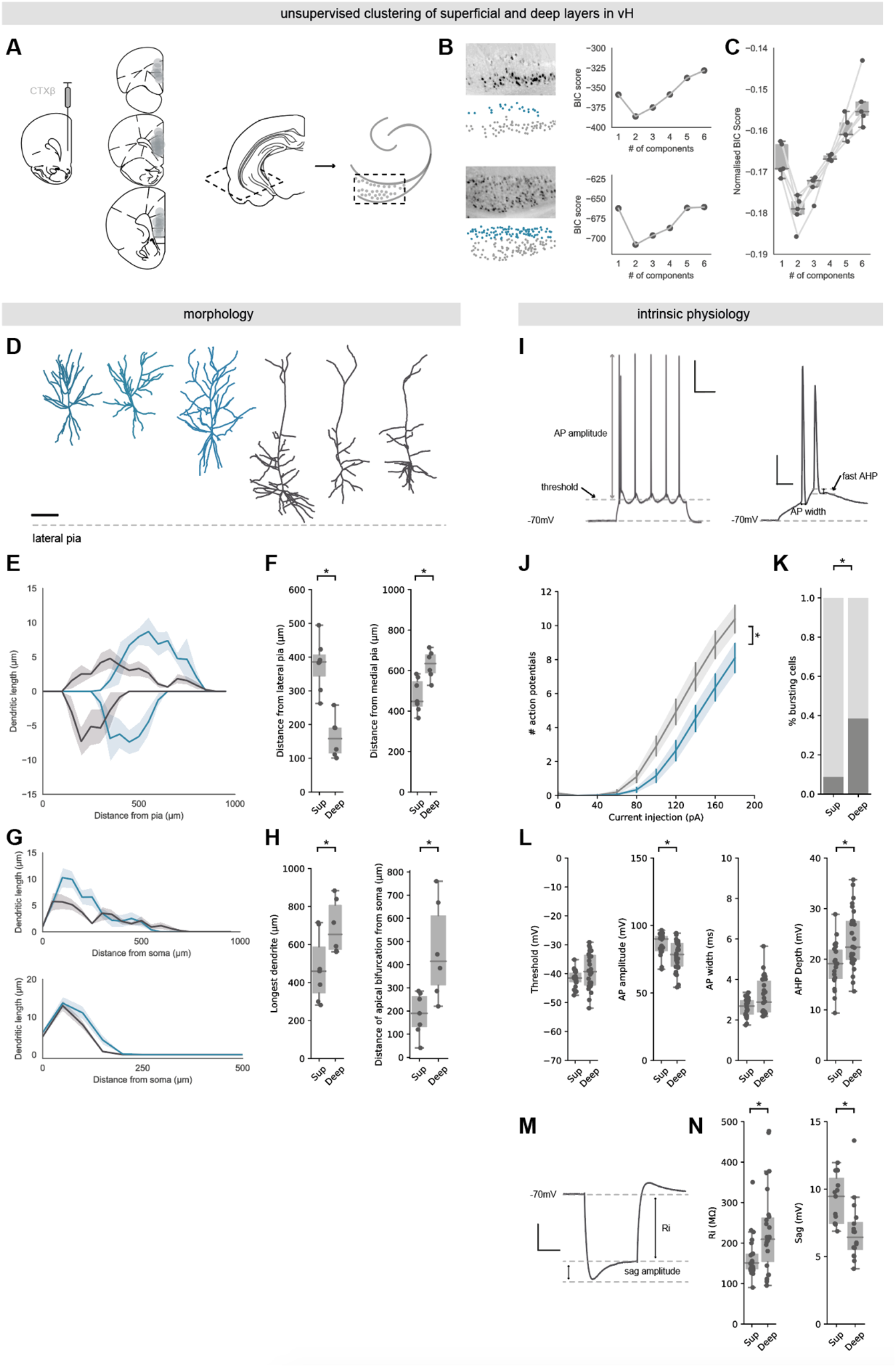
vH-PFC neurons form two morphologically and physiologically distinct populations segregated across the radial axis. **A)** *Left*, schematic of CTXβ injection into PFC and location of injections. Each injection is represented as a transparent fill. Thus, higher intensity reflects consistent labelling across injections. *Right*, Schematic showing orientation of transverse slice displayed in (**B**) and throughout manuscript. **B)** *Left*, two examples of CTXβ labelled cells in vH, one with clearly defined layered structure (top), and one where layering is less clear (bottom). *Right*, Bayesian information criterion (BIC) for gaussian mixture models with different numbers of components for neurons in each image. vH-PFC neurons are consistently best fit by models with two components, and these components are consistently split across the radial axis – one superficial and one deep (color coded in blue and grey underneath images on left). **C)** Summary of BIC scored across 7 injections showing consistency of two component fit. **D)** Example reconstructions of superficial (blue) and deep (dark grey) PFC-projecting hippocampal neurons. Scale bar = 100 μm, dotted line represents lateral pia (nearest cortex). **E)** Sholl analysis showing average dendritic length of apical (top) and basal (bottom) dendrites with increasing distance from the lateral pia for superficial (blue) and deep (grey) layer vH-PFC neurons. Each population samples distinct range of the radial hippocampal axis. **F)** Quantification of distance of the soma of recorded neurons in deep and superficial layers from the lateral pia (towards cortex - *left*), and medial pia (towards dentate gyrus – *right*). **G)** Sholl analysis of apical (top) or basal (bottom) dendrites with increasing distance from the soma for superficial (blue) and deep (grey) layer vH-PFC neurons. Superficial neurons have more complex dendritic trees. **H)** *Right*, quantification of the distance of the apical bifurcation from the soma in superficial and deep neurons. *Left*, quantification of the distance from the farthest dendrite tip to the soma. Deep layer neurons have longer apical dendrites. **I)** Example voltage traces from a deep layer PFC-projecting cell in response to current injections (140 pA). Trace on right is a zoom in of the first two events of the first trace. Arrows point at different aspects of the traces analyzed below. Scale bars = 100 ms, 20 mV; 10 ms, 20 mV. **J)** Quantification of number of action potentials elicited by somatic current injection in superficial (blue) and deep (black) neurons. **K)** Quantification of the proportion of bursting neurons after positive current injections. **L)** Quantification of (left to right) action potential (AP) threshold, AP amplitude, AP half-width, and depth of the fast after hyperpolarization (AHP). **M)** Example voltage trace from a deep layer PFC-projecting cell in response to current injections (− 160 pA). Arrows point at different aspects of the traces analyzed below. Sacle bar = 200 ms, 10 mV. **N)** Quantification of input resisteance and sag amplitude after negative current injections.

**Sup. Fig 2.**
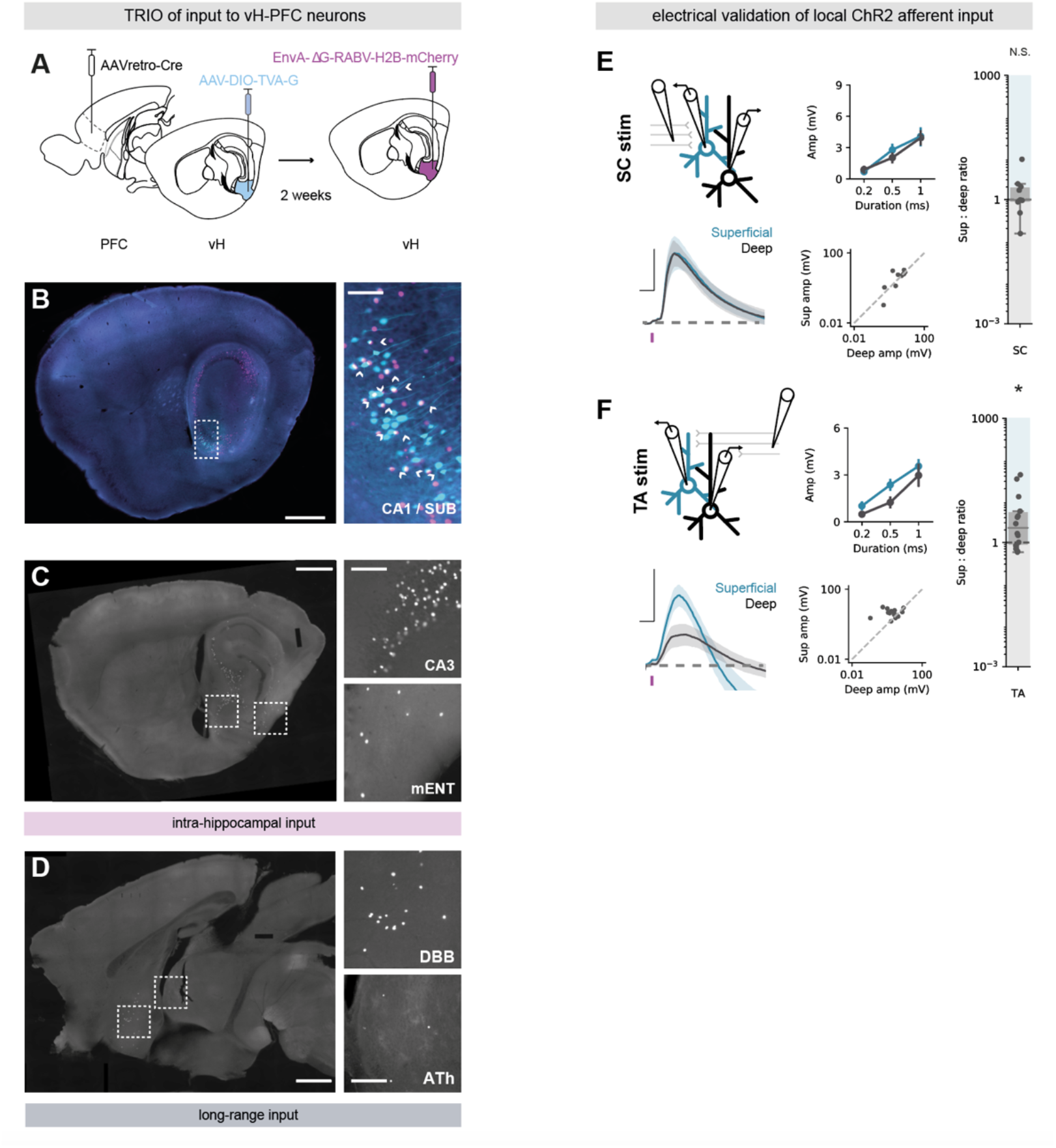
Quantification of TRIO-labelled inputs and electrically stimulated validation of CA3 and ENT input. **A)** Schematic of TRIO injection strategy. AAVretro-CAG-Cre was injected in PFC, and rabies helper proteins were injected into vH to limit subsequent rabies infection to hippocampal PFC projection neurons. 2 weeks later, EnvA pseudotyped rabies was injected in vH to label presynaptic neurons that synapse onto PFC-projecting hippocampal neurons with nuclear-localized mCherry. **B)** *Left*, Injection site in a sagittal section showing TVA and G protein expressing hippocampal neurons (cyan), and rabies labelled neurons (magenta). Scale bar = 1000 μm. *Right*, Zoom in to white dotted box. Co-labelled neurons represent starter neurons (arrows). Scale bar = 100 μm. **C)** Sagittal section showing insets of rabies-labelled cells in intra-hippocampal input regions CA3 and mENT. Scale bar = 1000 μm, 200 μm. **D)** Sagittal section showing insets of rabies-labelled cells in extra-hippocampal input regions DBB and ATh. Scale bar = 1000μm, 200 μm **E)** *Top left*, Schematic showing experimental setup. Retrobeads were injected into PFC. 2 weeks later Schafer collaterals (SC) were electrically stimulated to mimic CA3 activity, and connectivity was assessed using paired recordings of superficial and deep vH-PFC neurons in acute slices. *Bottom left*, Average stimulation-evoked responses in superficial (blue) and deep (black) layer PFC-projecting hippocampal neurons in response to cCA3 input. Scale bar = 10 ms, 2 mV. Purple tick represents the time of stimulus. *Middle*, summary of amplitude of Sup and Deep responses to increasing stimulus intensity (top), and amplitudes of individual pairs at 0.5 ms duration (bottom). *Right*, summary of the ratio of superficial : deep neuron EPSP. Higher values mean input is biased to superficial neurons, low values towards deep layer neurons. Note log scale. CA3 input is equivalent onto superficial and deep layer neurons. **F)** As in (**E**) but for stimulation of temperoammonic axons to mimic ENT activity. Scale bar = 10 ms, 2 mV. ENT input is biased towards activation of superficial layer neurons.

**Sup. Fig 3.**
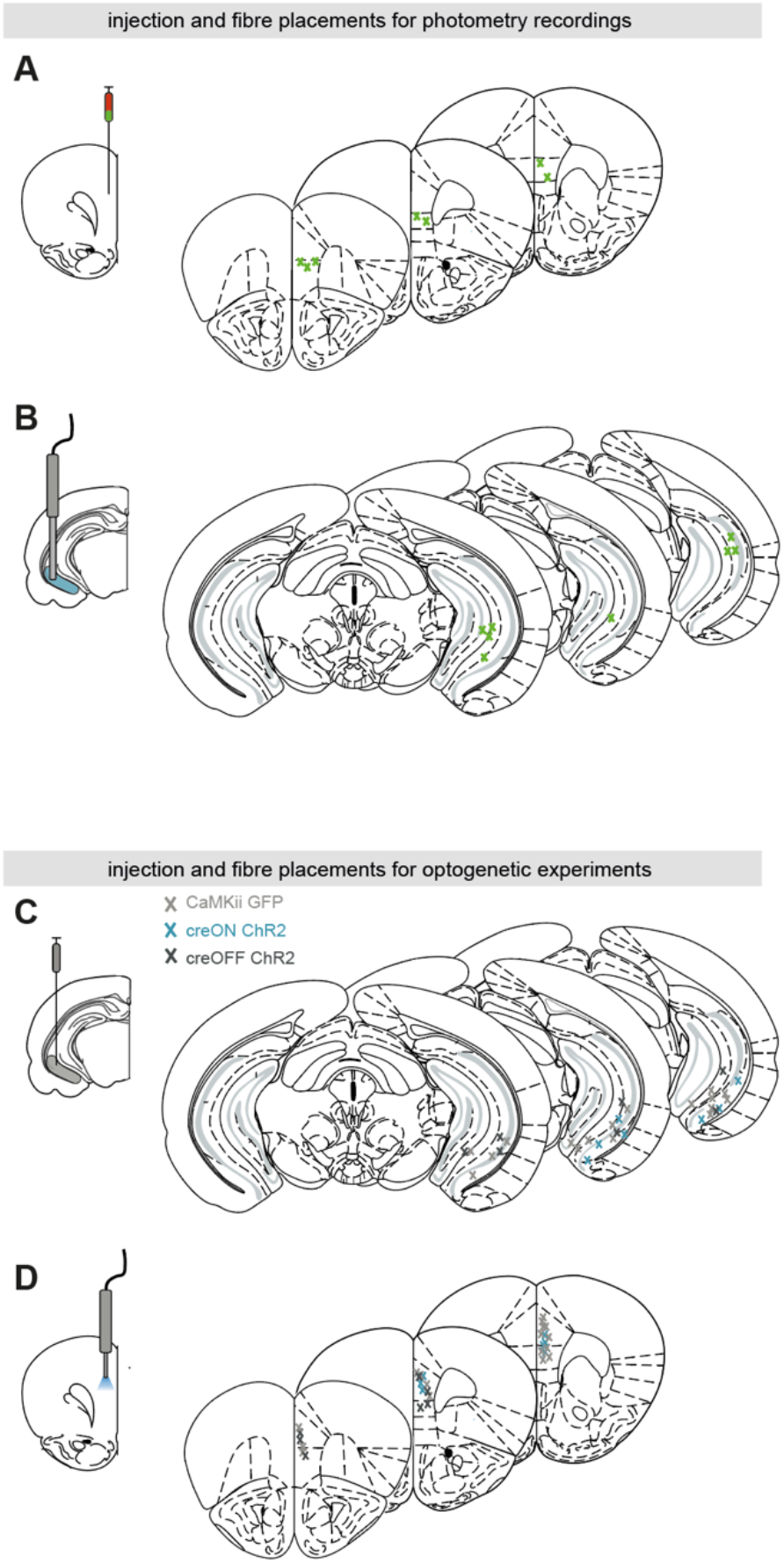
Injection site and fiber placements for in-vivo calcium imaging and optogenetics. **A)** *Left*, Schematic of viral injection of a 50:50 mix of AAVretro RGeCO1a and AAVretro GCaMP6f into PFC. Right, histology showing the location of the injection sites across all mice. **B)** as in (**A**) but for location of imaging fiber in vH. **C-D**) As in (**A,B**) but for injections of creON ChR2, creOFF ChR2 and GFP into vH, and fiber implants into PFC.

## SUPPLEMENTARY STATISTICAL TEST SUMMARY

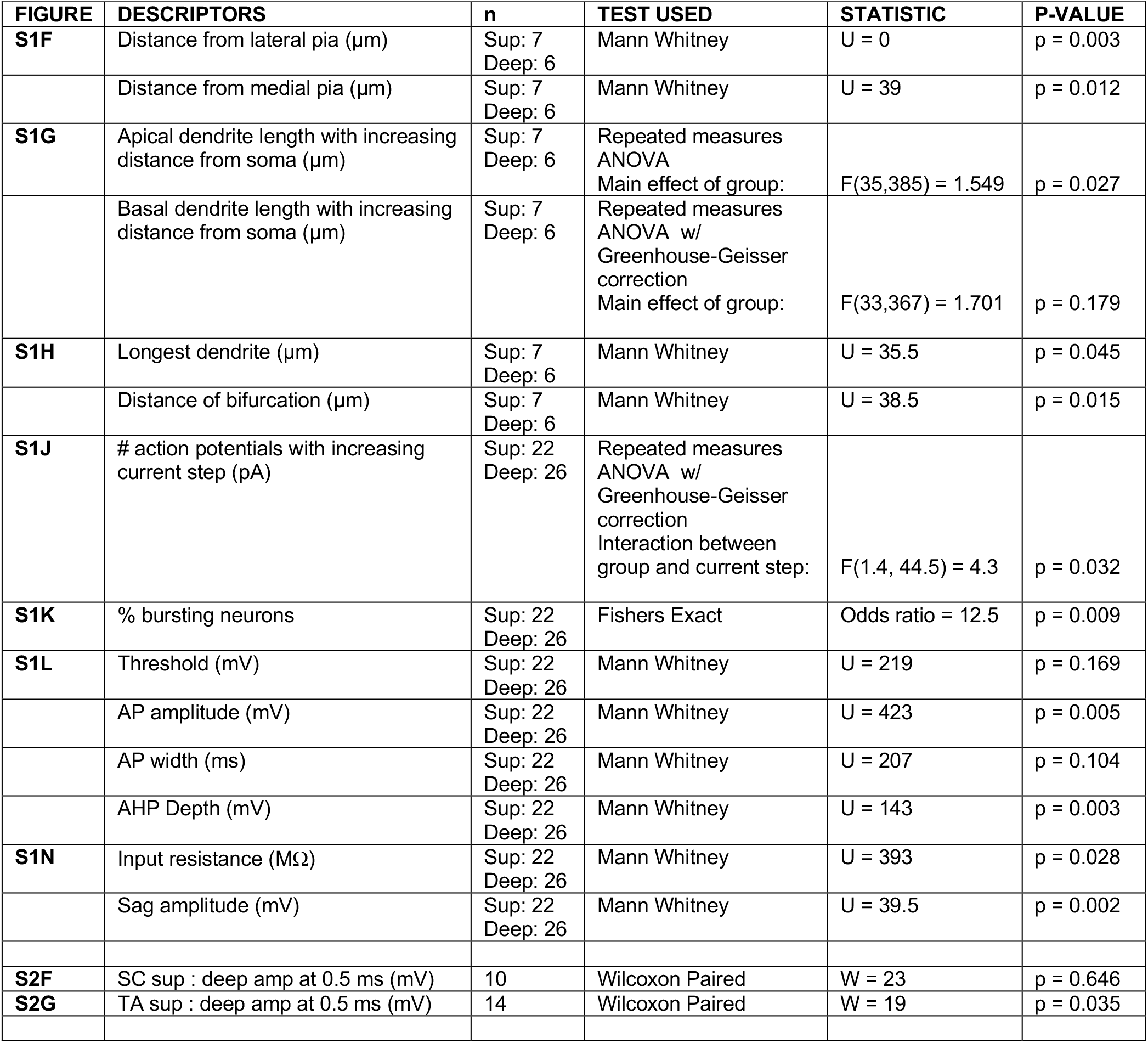

| FIGURE | DESCRIPTORS | n | TEST USED | STATISTIC | P-VALUE |
| --- | --- | --- | --- | --- | --- |
| **S1F** | Distance from lateral pia ($\mu\text{m}$) | Sup: 7<br>Deep: 6 | Mann Whitney | U = 0 | p = 0.003 |
| | Distance from medial pia ($\mu\text{m}$) | Sup: 7<br>Deep: 6 | Mann Whitney | U = 39 | p = 0.012 |
| **S1G** | Apical dendrite length with increasing distance from soma ($\mu\text{m}$) | Sup: 7<br>Deep: 6 | Repeated measures ANOVA<br>Main effect of group: | F(35,385) = 1.549 | p = 0.027 |
| | Basal dendrite length with increasing distance from soma ($\mu\text{m}$) | Sup: 7<br>Deep: 6 | Repeated measures ANOVA w/<br>Greenhouse-Geisser correction<br>Main effect of group: | F(33,367) = 1.701 | p = 0.179 |
| **S1H** | Longest dendrite ($\mu\text{m}$) | Sup: 7<br>Deep: 6 | Mann Whitney | U = 35.5 | p = 0.045 |
| | Distance of bifurcation ($\mu\text{m}$) | Sup: 7<br>Deep: 6 | Mann Whitney | U = 38.5 | p = 0.015 |
| **S1J** | # action potentials with increasing current step (pA) | Sup: 22<br>Deep: 26 | Repeated measures ANOVA w/<br>Greenhouse-Geisser correction<br>Interaction between group and current step: | F(1.4, 44.5) = 4.3 | p = 0.032 |
| **S1K** | % bursting neurons | Sup: 22<br>Deep: 26 | Fishers Exact | Odds ratio = 12.5 | p = 0.009 |
| **S1L** | Threshold (mV) | Sup: 22<br>Deep: 26 | Mann Whitney | U = 219 | p = 0.169 |
|  | AP amplitude (mV) | Sup: 22<br>Deep: 26 | Mann Whitney | U = 423 | p = 0.005 |
|  | AP width (ms) | Sup: 22<br>Deep: 26 | Mann Whitney | U = 207 | p = 0.104 |
|  | AHP Depth (mV) | Sup: 22<br>Deep: 26 | Mann Whitney | U = 143 | p = 0.003 |
| **S1N** | Input resistance ($\text{M}\Omega$) | Sup: 22<br>Deep: 26 | Mann Whitney | U = 393 | p = 0.028 |
|  | Sag amplitude (mV) | Sup: 22<br>Deep: 26 | Mann Whitney | U = 39.5 | p = 0.002 |
| **S2F** | SC sup : deep amp at 0.5 ms (mV) | 10 | Wilcoxon Paired | W = 23 | p = 0.646 |
| **S2G** | TA sup : deep amp at 0.5 ms (mV) | 14 | Wilcoxon Paired | W = 19 | p = 0.035 |

## Notes

### Competing Interest Statement

The authors have declared no competing interest.

